# Natural Selection as the Sum over all Histories

**DOI:** 10.1101/2021.02.24.432748

**Authors:** C E Neal-Sturgess

**Affiliations:** Emeritus Professor of Mechanical Engineering, University of Birmingham

## Abstract

As evolution can be connected to the principle of least action, and if it is depicted in evolution-space versus time then it corresponds to the direction of ultimate causation. As an organism evolves and follows a path of proximate causation, if the vector is closely parallel to that of the Ultimate Causation then the changes will confer desirable attributes which will lead to further development. If, however, the variations do not occur in a direction close to that of the ultimate causation vector the evolved organism will quickly die out. Therefore Natural Selection may be viewed as similar to Feynman’s “sum over all histories”. This approach is compatible with both Neutral Theory and Selection, as it includes both positive and negative mutations and selection. Therefore, the principle of least action gives a direction, but not a purpose, to evolution.

## Background

Since the publication of “On the Origin of Species by Natural Selection” by Charles Darwin in 1859, there have been a number of attempts to link it to other scientific principles, notably the Principle of stationary Action, known popularly as “Least Action” [Nahin 2004]. In their paper Natural selection for Least Action (Kaila and Annila 2008) they depict evolution as a process conforming to the Principle of Least Action (PLA). This paper, although not giving any experimental evidence, shows that evolution, if conforming to the Second Law of Thermodynamics, will follow a trajectory of maximum Entropy production, which conforms to the PLA. To demonstrate this they rewrote the Gibbs-Duhem relationship in terms of possible states, to give a differential equation of evolution. This is a convincing argument as biology at root is a physical process, as expounded by Schrodinger 1944, and all physical processes are governed by the Second Law of thermodynamics.

The PLA first arose, in modern times in the 17^th^ century, and was popularised by Maupertius [Terrall M 2002] although the major rigorous work was done by Euler [Kaila & Annila 2008]. The PLA was Feynman’s guiding principle throughout this life [Feynman 1985]. As Wittgenstein 1922 said “people were using the Principle of Least Action before they knew it existed”.

The Principle of Least (Stationary) Action is stated simply as:

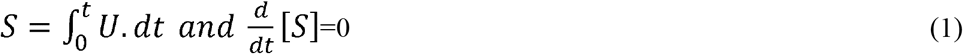

Where U is the Gibb’s Free Energy.

The second law can be stated as:

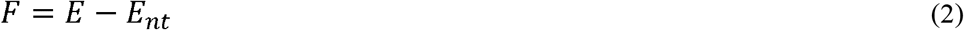

Where: E is the total energy of the body, F is the “Gibbs Free Energy” and E_nt_ is the entropy.

## Results

Here it is hypothesised that an analogue exists to Feynman’s “Sum Over All Histories” [Feynman 1985]. Richard Feynman, Nobel Laureate, was fascinated by the PLA for most of his life, and it led to one of his greatest breakthroughs in Quantum Electrodynamics or QED – the quantum theory of light [Feynman 1985]. Feynman considered that a Photon traversing a path from A to B in space-time was free to traverse any path. However, when the sum of all paths is taken, because of phase differences, the most likely path is that conforming to the PLA; this is known as the “Sum Over All Histories”.

It is assumed here that there is a general overall direction of evolution. The vector of evolution is called here the Ultimate Causal Direction (green vector), in accordance with the terminology used in evolutionary biology(Mayr 2001), and this should conform to the PLA [Kaila & Annila 2008], in conformity with the external conditions, and could be shown graphically as shown in Fig.1.

**Fig.1.**
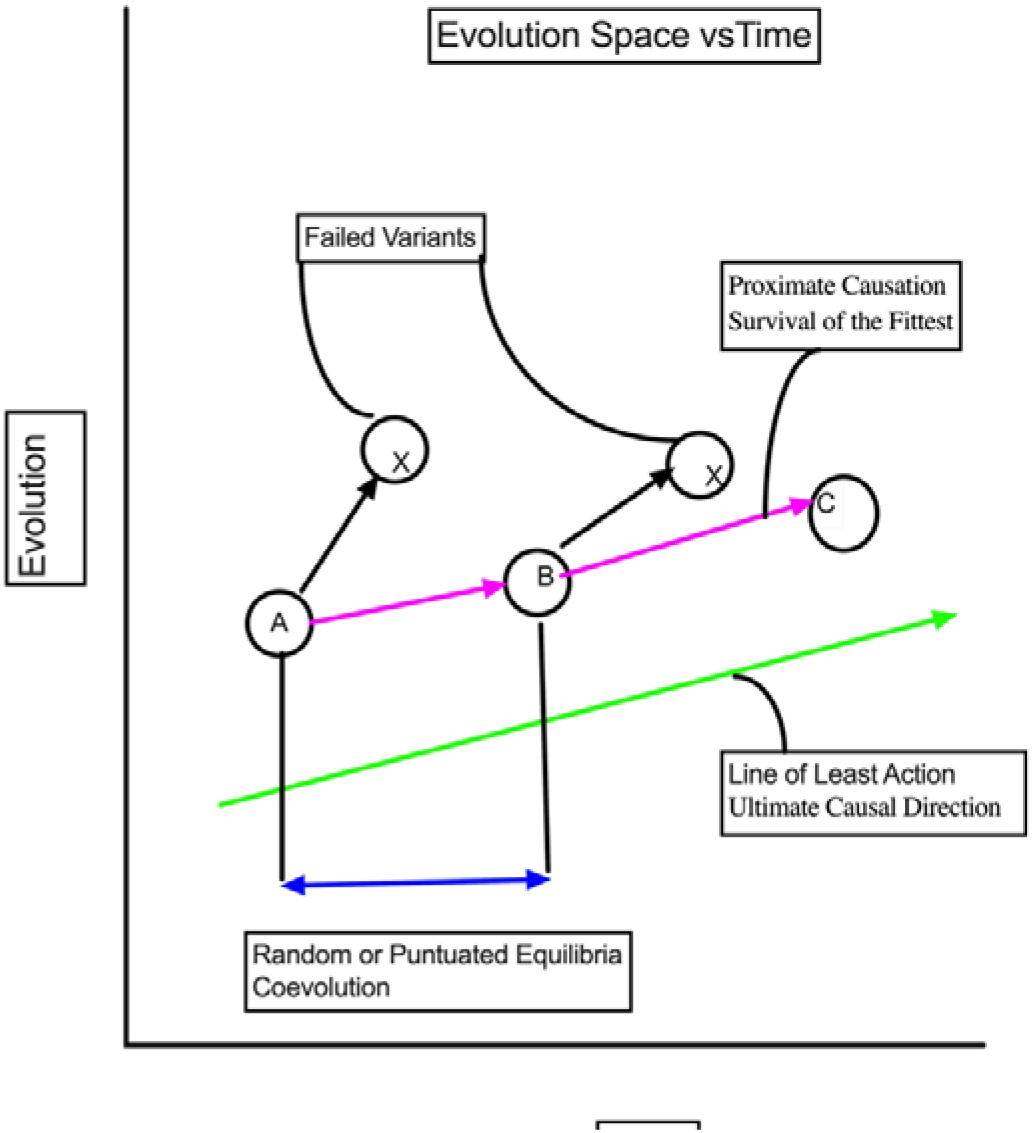
Evolution-Space vs Time.

If during any genetic mutation leading to a change in the organism takes place (event A) this will in general be random, under constraints where the genome stays constant but the phenotype changes [Holmes 2013] and confers on the next generation of the organism’s attributes which may or may not lead to a greater probability of survival in the external climate [Loewe 2010]. These changes to the phenotype are in evolutionary biological terms called “Proximate Causation”. It is proposed that Proximate Causation will lead to greater survivability if it is parallel, or closely parallel, to the direction of PLA, for the external conditions at the time (fitness). This is shown in Fig.1. as the red vector A-B-C. In this diagram, those variants not reasonably parallel to the trajectory of ultimate causality, the black lines ending in blobs, will die out. Further Epigenetic changes [Dupont 2006] could also more closely align the red vector with the Ultimate Causality. Although this diagram as an axis of Time, the interval between the changes is not known, as shown by the caption. The eventual unfolding of Proximate Causation towards Ultimate Causation as determined by both thermodynamics and the environment can be considered as analogous to Feynman’s “Sum over all Histories”, as the mutations leading to Proximate causation may be viewed as exploring all the possible states in evolution space, but only those approximately parallel to the environment vector will survive to generate new variants and lead to the Ultimate Causality; the other variants will die out [Loewe 2010]. Hence evolution towards the Ultimate Causality can be seen as the ultimate goal of the sum over all proximate evolutionary histories and corresponds to Natural Selection. This approach is compatible with both Neutral Theory [Kimura 1983, Durant 2008] and Selection, as it includes both positive and negative mutations.

This is not to say the evolution to the PLA is teleological, there is no “purpose”. It is simply that those evolutionary lines arising from proximate causality that are closely parallel to the line of PLA, are best adapted to whatever external environments that exist at the time, and so survive longer, and have a greater probability of reproduction (heritable); therefore, evolution has a direction but not a purpose.

There are also interesting developments in evolutionary thinking where some are beginning to posit that organisms make changes to their phenotype, as opposed to their genotype, by adjusting to their environment (Holmes 2010). This tendency here would be to add a rotation to the Proximate Causality, hence bringing the vector closer to the direction of the Ultimate Causality, in the direction of PLA.

Most investigators consider that natural selection is the cumulative effects of the genetic mutations, which is the position adopted here. Other investigators however classify natural selection as one of the mechanisms of mutation [Sanjuan & Domingo-Calap 2016], which is not adopted here.

However, there is a measurement problem in evolution through the fossil record in that the rate of progress is inversely proportional to the timescale over which it is measured, leading to Gould’s “Punctuated Equilibrium” ideas (Eldridge & Gould 1972, Gould 2002). Furthermore, another measurement problem is that of actual findings and observation. In general, the probability of finding a fossil will be proportional to the number of the species that existed, and so the time the fossil type lived. Therefore, the probability of finding a fossil type is dependent on how successful that fossil type was, and so most of the changes that have occurred in a species will probably never be found because they died out quickly. This is an exceptionally complicated problem, and as said by [Loewe and Hill] that it is “like looking for a needle in a haystack’; however, advances in Evolutionary Genetics may give answers in the future.

## Conclusions

Evolution can be connected to the Principle of Least Action, and if it is depicted in “Evolution-Space vs Time” then it corresponds to the direction of Ultimate Causation. As an organism evolves and follows a path of Proximate Causation, if the vector is closely parallel to that of the Ultimate Causation then the changes will confer desirable heritable attributes which will lead to further development. If, however the variations do not occur in a direction close to parallel to the of the Ultimate Causation vector the evolved organism will quickly die out. Therefore Natural Selection may be viewed as similar to Feynman’s “sum over all histories”. This approach is compatible with both Neutral Theory and Selection, as it includes both positive and negative mutations and selection. Therefore, the Principle of Least Action gives a direction, but not a purpose, to Evolution.

## Notes

### Competing Interest Statement

The authors have declared no competing interest.

### Summary of Updates

This revision is to separate this topic from the differential equation contained in the first version.

